# *Escherichia coli* Lrp regulates one-third of the genome via direct, cooperative, and indirect routes

**DOI:** 10.1101/276808

**Authors:** Grace M. Kroner, Michael B. Wolfe, Peter L. Freddolino

**Author notes:** GMK and MBW contributed equally.

## Abstract

The global regulator Lrp plays a crucial role in regulating metabolism, virulence and motility in response to environmental conditions. Lrp has previously been shown to activate or repress approximately 10% of genes in *Escherichia coli*. However, the full spectrum of targets, and how Lrp acts to regulate them, has stymied earlier study. We have combined matched ChIP-seq and RNA sequencing under nine physiological conditions to map the binding and regulatory activity of Lrp as it directs responses to nutrient abundance. In addition to identifying hundreds of novel Lrp targets, we observe two new global trends: first, that Lrp will often bind to promoters in a poised position under conditions when it has no regulatory activity, and second, that nutrient levels induce a global shift in the equilibrium between non-specific and sequence-specific DNA binding. The overall regulatory behavior of Lrp, which as we now show regulates 35% of *E. coli* genes directly or indirectly under at least one condition, thus arises from the interaction between changes in Lrp binding specificity and cooperative action with other regulators.

## Introduction

Regulation in response to changing nutrient conditions is a vital characteristic for free-living microbes, which must rapidly sense and respond to their environment in order to optimize fitness. The frequently studied model microbe *Escherichia coli* (*E.coli*) uses a hierarchical regulatory architecture to coordinate responses to environmental changes, with the activity and actions of dozens of specific transcription factors organized by seven global regulators: ArcA, FNR, Fis, CRP, IHF, H-NS and Lrp [1]. *E. coli* Lrp is the eponymous member of the Lrp/AsnC protein family, and regulates 70% of the 215 genes with differential expression upon entrance to stationary phase [2]. It influences a variety of cellular processes: amino acid synthesis, degradation and transport, porin expression, and pilus formation [3,4]. The latter represents an example of how Lrp homologues have recently been tied to expression of virulence genes [5–10].

Lrp itself is an 18 kD protein containing a helix-turn-helix DNA binding domain and a regulator of amino acid metabolism (RAM) domain [11]. *In vivo*, it is thought to exist in an equilibrium between octameric and hexadecameric states [12]. Binding of leucine to the RAM domain is known to favor formation of octamers over hexadecamers [13] and to increase the nonspecific DNA binding affinity of Lrp [14]. In addition, the presence of leucine affects Lrp’s regulatory role. Depending on the target, Lrp either activates or represses transcription, and in turn, leucine binding to Lrp either potentiates or inhibits Lrp function [15]. Recent studies also indicate that Lrp may respond to other amino acids, including alanine, methionine, isoleucine, histidine and threonine [16]. Cho *et al*. [15] performed chromatin-immunoprecipitation (ChIP) using epitope-tagged Lrp under three conditions, resulting in some expansion of the known Lrp regulon. However, in comparison to other global regulators, the Lrp regulon as currently known is relatively small, suggesting that all targets have not been identified. In addition, although the concentration of Lrp is not as high as some nucleoid-associated proteins like H-NS and HU, it is expressed to a similar degree as CRP [17]. Thus one might expect that their regulons would be of similar size; however, there is currently a dramatic discrepancy between these proteins with CRP annotated as regulating 572 genes, while Lrp only regulates 109 [18]. Based on estimates about the levels of Lrp and the percentage found free of the nucleoid [14], we estimate that there should be between 400 and 500 Lrp octamers bound and capable of modulating transcription levels under logarithmic growth in both rich and minimal media conditions. Additionally, we still lack a mechanistic understanding of how Lrp regulation occurs.

Making use of a carefully refined ChIP-grade antibody for Lrp, we employed chromatin-immunoprecipitation followed by DNA sequencing (ChIP-seq) of native Lrp in a variety of media conditions and growth phases to assess the full spectrum of Lrp binding sites. Coupled RNA-seq experiments on both wild type (WT) and Lrp knockout (*lrp::kanR*) cells enabled us to distinguish between productive and apparently non-functional binding events, and between direct and indirect Lrp regulatory targets. This rich, high-confidence data set has allowed us to categorize hundreds of novel direct and indirect Lrp targets, expanding Lrp’s regulon to 35% of genes in *E. coli* (roughly one-fifth of which are direct targets of Lrp), compared to the 10% previously documented. In addition, we identify a surprising but highly prevalent mode of Lrp binding in which Lrp binds to a site under many physiological conditions, but only alters transcription under certain conditions, similar to poised transcription factor binding in eukaryotes [19,20]. We show that some of Lrp’s poised regulation may be explained by interactions with other regulatory factors such as the nitrogen-response sigma factor, σ^54^. Despite extensive efforts, we were unable to identify systematically enriched sequence determinants sufficient to either explain transitions from poised to active regulation, or predict Lrp activation from Lrp repression. However, we did observe a shift in Lrp’s DNA binding specificity in response to varying nutrient conditions. The conservation of Lrp across many species of bacteria and archaea [21] argues for its critical role in organismal survival, and here we provide the most comprehensive picture of the Lrp regulon in *E. coli* to date, establishing rules for Lrp behaviour that will likely illuminate study of the protein in many species. The general principles of Lrp’s behavior across conditions may also serve as a template for other bacterial global regulators.

## RESULTS

### ChIP-seq and RNA-seq identify hundreds of novel Lrp targets

We performed both ChIP-seq and RNA-seq on WT and Lrp knockout (*lrp::kanR*) cells to establish a global picture of Lrp binding and regulatory effects in nine physiological conditions. Conditions and time points will be referenced as follows: the time points are denoted X_Log (logarithmic phase), X_Trans (transition point), and X_Stat (stationary phase), where the X may be MIN (minimal media), LIV (minimal media supplemented with branched-chain amino acids), or RDM (rich defined media). Overall, the combination of Lrp binding data from the ChIP-seq experiments and the expression data from the RNA-seq experiments resulted in identification of hundreds of novel Lrp targets. We document a ten-fold range (between 65 and 668) in the number of Lrp peaks identified across the nine physiological conditions examined here. Fewer Lrp binding sites are identified in media with higher nutrient conditions (either LIV or RDM) relative to the MIN (Fig 1A), in agreement with previously published Lrp ChIP data [15] and with Lrp’s known role as a regulator which responds to decreasing nutrient levels. However, our data identifies between two- and five-fold more binding sites overall than previous studies. In general, we document more Lrp binding sites at later time points (Trans and Stat) relative to Log (Fig 1A); again in agreement with previously published Lrp ChIP data [15] and with the known role of Lrp as being a critical regulator at the transition to stationary phase. Comparing our data to previously published ChIP-ChIP studies [15], we identify extensive overlap in binding locations: 96% of sites in prior ChIP-ChIP data are reproduced in our data at MIN_Log (26.8 fold enrichment; p < 0.001, permutation test, r=1000; here and throughout the manuscript we use r to refer to the number of replicates used for resampling tests), 44% at LIV_Log (121.7 fold enrichment, p < 0.001, permutation test, r=1000) and 84% at MIN_Stat (15.5 fold enrichment, p < 0.001, permutation test, r=1000). Comparing at the level of genes which are identified as having a Lrp-dependent change in expression as measured by RNA-seq, our data set overlaps with 78% of the known targets in RegulonDB (1.50 fold enrichment, p < 0.001, permutation test, r=1000), 77% of the previously identified ChIP-ChIP targets (1.49 fold enrichment, p < 0.001, permutation test, r=1000) [15], and 89% of the previously identified microarray targets (1.72 fold enrichment, p < 0.001, permutation test, r=1000) [2], showing good agreement across the variety of strains and media conditions present in the compared studies, despite some variations in precise experimental conditions. We also identify over 900 novel Lrp binding sites, and 2104 genes with previously undocumented Lrp-dependent expression.

**Figure 1:**
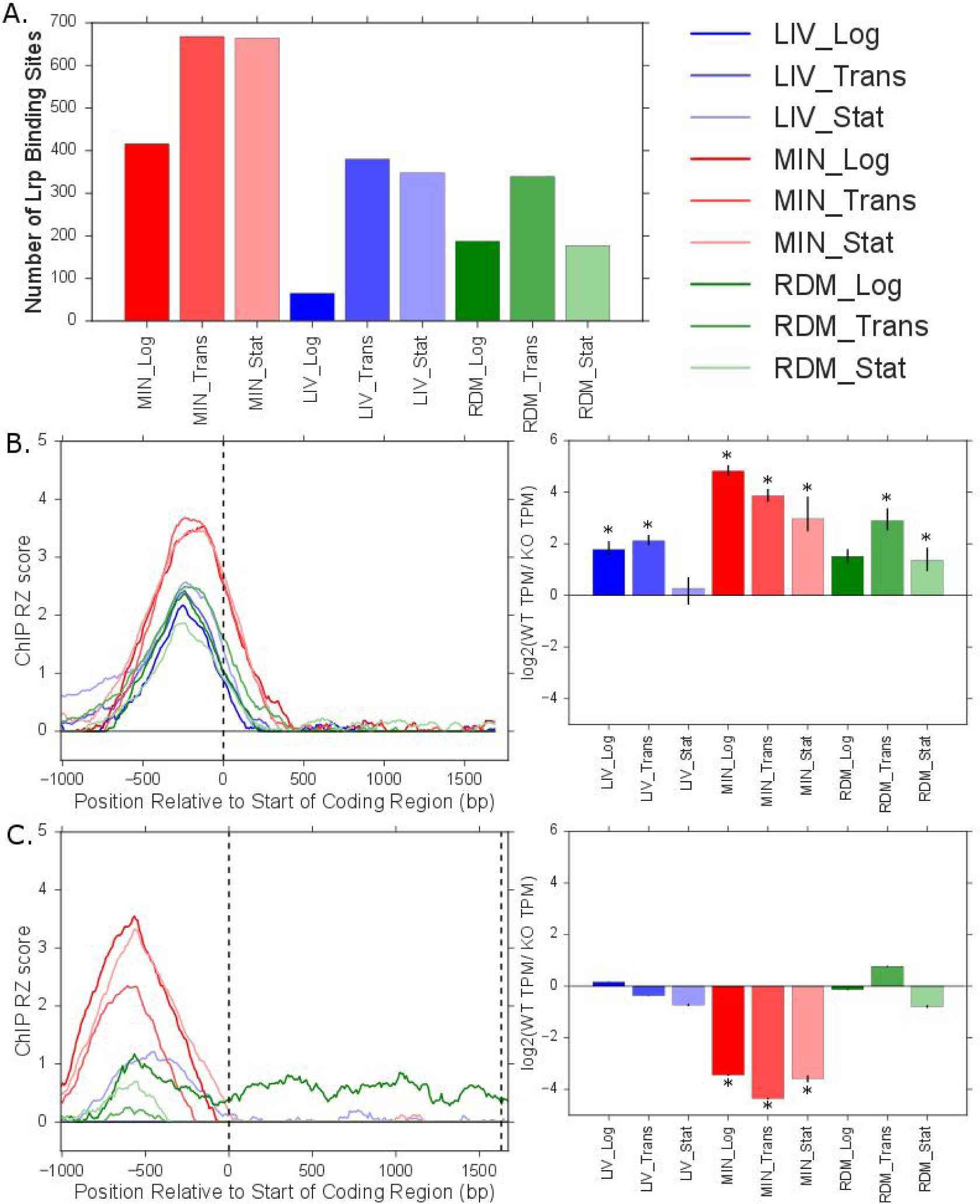
*ChIP-seq data shows agreement with previous data and reveals novel Lrp binding sites.* **A**. Total number of non-overlapping high-confidence Lrp binding sites identified in each condition. **B.** ChIP robust Z-score (left) and RNA-seq expression change (log2(WT/KO); right) for known Lrp activated target *ilvI*. Dashed vertical lines on the ChIP robust Z-score graph mark the start and end of the gene coding region. Error bars for the RNA-seq data indicate a percentile based 95 % confidence interval from 100 bootstrap replicates of TPM estimates. Stars indicate a significant difference in RNA abundance between WT and *lrp::kanR* strains (Wald Test qvalue of < 0.05 and a genotype log fold change coefficient magnitude of > 0.5; see Methods for details). **C.** ChIP robust Z-score (left) and RNA-seq expression change (log2(WT/KO); right) for known Lrp repressed target *oppA,* panels as in **B**.

Many well-studied Lrp targets are reproduced in our data. IlvI (b0077) is an enzyme critical for valine and isoleucine biosynthesis that is known to be activated by Lrp [22]. Consistent with prior work, we see a strong Lrp binding signal at the *ilvI* transcription start site (Fig 1B, left panel), and a Lrp-dependent activation of *ilvI* transcription in several media conditions (Fig 1B, right panel). The extent of activation is weakened or eliminated completely in LIV or RDM conditions, in agreement with previous studies showing that leucine inhibits the Lrp-mediated activation of *ilvI* [23].

A strong Lrp binding signal under MIN conditions is also evident at the promoter region for OppA, a protein critical for oligopeptide transport [24]. Lrp is known to repress expression of the *oppABCDF* operon in the absence of leucine [25]. Accordingly, we see Lrp-dependent repression of *oppA* under MIN conditions (Fig 1C, right panel). The Lrp binding signal is strongly attenuated, and there are no Lrp-dependent expression effects, during LIV and RDM conditions (Fig 1C).

### Global analysis reveals that Lrp has condition-specific modes of binding and regulation

Global regulators are known to act both directly, by binding target sites and modulating transcription levels, and indirectly, by modulating the expression of transcription factors which have their own targets [1]. Previously, most focus on Lrp regulation has been at the direct target level. By comparing the binding data from our ChIP-seq experiments and the corresponding expression data provided by our RNA-seq experiments, we are able to identify and categorize both direct and indirect targets under a variety of physiological conditions. Direct and indirect targets are both characterized by Lrp-dependent changes in expression, but only direct targets have a Lrp binding signal in their regulatory regions, defined as 500 bp upstream and downstream of the annotated transcription start site (Fig 2A). In addition, our data shows many examples of a converse mode of Lrp activity, in which binding of Lrp is apparent at a particular promoter, but there are no Lrp-dependent changes in expression (these sites comprise 65-94% of all instances of Lrp binding across the conditions that we studied). We refer to such cases as instances of nucleoid-associated protein (NAP) activity of Lrp, thus described due to the similarity of Lrp’s behavior at these locations with highly abundant,low-specificity NAPs such as H-NS and HU [26]. In our data set, neither NAP-type Lrp targets nor genes unconnected to Lrp show Lrp-dependent RNA expression changes by definition, but the NAP targets have a Lrp binding signal (Fig 2A). This is apparent, for example, at the *ybjN* gene; its promoter region is always bound by Lrp, but never exhibits a significant change in expression, thus making it a NAP-type target under all conditions (Fig 2B). Interestingly, YbjN is proposed to play a role in stress response and motility [27], areas to which many of Lrp’s targets are known to belong. The consistent binding of *ybjN*’s promoter by Lrp coupled with the similarity of its role to other Lrp targets suggests that Lrp may always be poised to regulate YbjN, and that it may be a direct target of Lrp under conditions not tested here.

**Figure 2:**
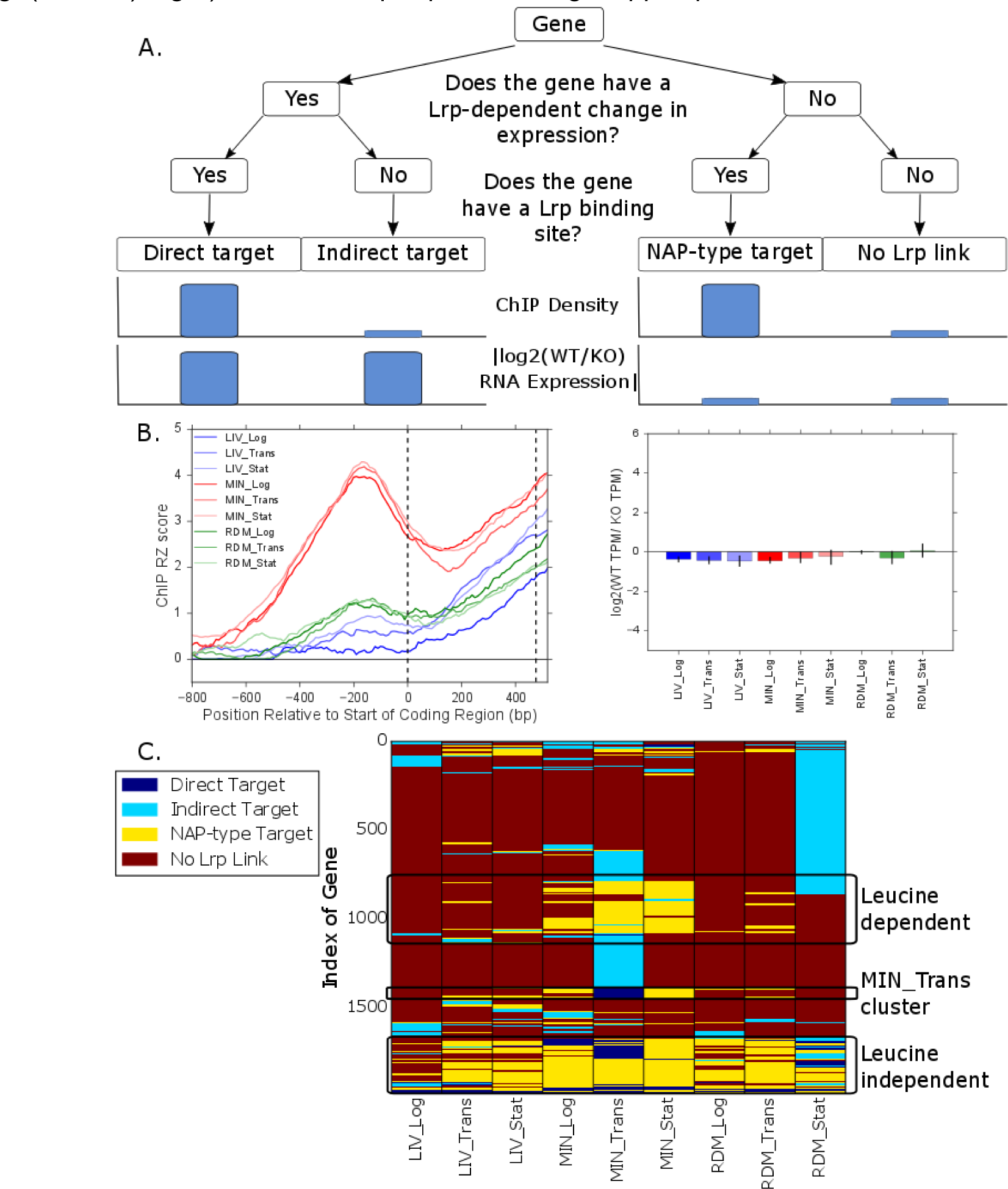
*Lrp regulates genes both directly and indirectly.* **A.** Schematic showing how genes were categorized: direct targets of Lrp (Lrp-bound regulatory region and with a significant RNA expression change between WT and *lrp::kanR* cells), indirect targets (not bound but with a significant RNA expression change), NAP targets (bound but with no significant RNA expression change), or not linked (not bound and no significant RNA expression change). Filtering was done independently for each condition. **B.** ChIP robust Z-score (left) and RNA-seq expression change (log2(WT/KO); right) for Lrp NAP-type target *ybjN* (as in Fig 1B). **C.** Heat map indicating how each gene was classified in the nine experimental conditions. Genes with no Lrp link in any condition were removed from visualization. Genes were hierarchically clustered using a Manhattan distance metric and average linkage clustering. Black boxes mark out notable clusters of genes: those with leucine-dependent or -independent binding and those that are direct targets only under MIN_Trans.

Based on our RNA-seq data, we find that 1.7% to 29% of all *E. coli* genes are regulated by Lrp in each condition (Table 1), equaling 2320 unique Lrp-regulated genes (50% of total *E. coli* genes). However, due to the presence of operons in *E. coli*, in the analysis below we only categorized genes (as direct, indirect or NAP-type targets) if a transcription start site exists in the PromoterSet dataset in RegulonDB version 9.4 within 500 base pairs upstream of the start of the coding region, resulting in categorization of 2875 genes out of the 4658 present in *E. coli* MG1655. From our analysis of that categorizable subset, we note that 35% of all *E. coli* genes are regulated by Lrp, either directly or indirectly, in at least one condition. Out of those, about 81% are regulated indirectly, 13% are regulated directly, and 6% are labelled as indirect and direct targets in different conditions. Due to the restriction on categorizing genes noted above, the counts given here are an underestimate. If we assume each transcription unit is fully transcribed and therefore assign the Lrp categorization of the first gene to each subsequent gene in the transcription unit, that increases the Lrp regulon to 49% of all *E. coli* genes (2289 genes/4658 total genes), with 16% of that total being direct targets, 78% being indirect targets and 6% being categorized as both in different conditions. This extended estimate matches the fraction of genes that show a Lrp-dependent change in expression as measured by RNA-seq. In addition, since some later genes within a transcriptional unit have independent transcription start sites, there exists the possibility of those genes being categorized as indirect targets because the operon itself is Lrp-controlled even if there is no Lrp peak near the later gene in the operon. Given that we do not know how often the overall operon start site is used compared to the internal start sites, we are not able to place those genes unambiguously as indirect targets. We calculated how many of these ambiguous indirect targets were present per condition and noted that it is a fairly small percentage of total indirect targets, ranging from 1.2% to 12.0% (with a median of 5.7%). These ambiguous targets comprise 44 total genes in *E. coli* (Table S1). Given the small size of this gene subset, we proceeded with the original categories and included the ambiguous indirect targets with the indirect targets in the analysis below.

**Table 1:** *Genes with significant Lrp-dependent changes in expression.* Percentage is out of the total number of genes in *E. coli* (4658).

| Condition | Total Genes Significantly Upregulated by Lrp | Total Genes Significantly Downregulated by Lrp |
| --- | --- | --- |
| MIN_Log | 164 (3.52%) | 162 (3.48%) |
| MIN_Trans | 423 (9.08%) | 590 (12.67%) |
| MIN_Stat | 63 (1.35%) | 63 (1.35%) |
| LIV_Log | 109 (2.34%) | 217 (4.66%) |
| LIV_Trans | 87 (1.87%) | 123 (2.64%) |
| LIV_Stat | 80 (1.72%) | 68 (1.46%) |
| RDM_Log | 36 (0.77%) | 84 (1.80%) |
| RDM_Trans | 58 (1.25%) | 21 (0.45%) |
| RDM_Stat | 728 (15.63%) | 622 (13.35%) |

We next used hierarchical clustering to order the categorized genes (those 2875 genes with an annotated transcription start site) by assessing how similar their categorization assignments were over the nine sampled physiological conditions (Fig 2C). We immediately noted that genes can transition between labels, e.g. from direct target to NAP, depending on the media condition and time point. As seen with the case of YbjN above, these findings suggest that Lrp is often poised at a particular gene under many conditions, but must act combinatorially with some other factor or environmental stimulus in order to actually alter expression. In addition, we see evidence for leucine-independent and leucine-dependent binding; some genes are always NAP-type or direct targets (i.e. Lrp bound) regardless of condition (the leucine-independent group) and some are only bound during MIN conditions (the leucine-dependent group). There is no obvious cluster that is bound only under conditions of high leucine levels. We also observe a dramatic increase in the number of indirect targets at MIN_Trans and RDM_Stat, going from 150 to 510 indirect targets between MIN_Log and MIN_Trans, and 34 to 920 indirect targets from RDM_Trans to RDM_Stat. These conditions likely represent points in growth at which Lrp’s regulatory activity is particularly important for fitness.

### Lrp binding is enriched among regulatory regions of the genome

As detailed in the Methods section, our process for categorizing genes as Lrp targets involved testing whether there was a called Lrp peak within 500 bp upstream or downstream of each annotated transcription start site (TSS) in the *E. coli* genome. If there were multiple annotated transcription start sites, we took 500 bp upstream of the most distal TSS (relative to the start of the gene itself) and 500 bp downstream of the most proximal TSS. We classified those approximately 1000 bp windows as regulatory regions, and tested whether Lrp binding was significantly enriched in those regions. Overall, 48% of the *E. coli* genome falls into these regulatory regions. However, we observe between 62% and 86% of Lrp peaks appearing in regulatory regions. A permutation test in which the same size and number of peaks were randomly shuffled across the genome indicated that there is significant enrichment for Lrp binding in regulatory regions (Table S2). This strongly supports Lrp’s role as a specific regulatory protein.

The Lrp peaks not in regulatory regions were distributed in gene coding regions, between genes in a transcription unit, or in truly intergenic regions at relative ratios similar to the proportion of those regions on a genome-wide scale (Table S3). We investigated whether any of those peaks might affect full transcription of an operon, hypothesizing that Lrp binding in the middle of an operon might block RNA polymerase. From the RNA-seq data, we identified any genes that showed a Lrp dependent change in expression, were not classified (and so did not have their own transcription start site), had a Lrp response that was different from the first gene in the corresponding transcriptional unit, and had a Lrp peak within 1000 bp upstream or downstream of the gene coding region. Due to incomplete annotation of the *E. coli* genome, some of these genes appear to be ones that should have a unique transcription start site based on visual inspection of the genomic context. However, for the remaining examples, we compared the RNA-seq coverage to the location of the peak as identified by the Lrp ChIP signal. As seen for the binding at *ilvI*, we again note that Lrp binding does not guarantee a regulatory effect. Genes that have a strong internal Lrp binding site under all conditions do not evince a Lrp dependent change in expression at all times, and Lrp binding sites within an operon do not, in general, appear to hamper transcription (Fig S1). These findings again suggest that Lrp regulation is often dependent on cooperative interaction with other regulatory factors.

### Direct Lrp targets explain the Lrp-dependent regulatory effect at some indirect targets

Given the high proportion of indirect Lrp targets, and especially the dramatic increase in the number of indirect targets at MIN_Trans and RDM_Stat, we investigated whether some of the expression changes of those indirect targets can be explained by the activity of direct Lrp targets at those time points. As Lrp is a global regulator, we expected to find that some percentage of the indirect targets could be explained by considering the known targets of the transcriptional regulators categorized as direct targets under that condition. We would expect that in such cases, we should observe an enrichment among Lrp indirect targets of genes known to be regulated by Lrp direct targets. We observe significant, albeit small, enrichment of explainable indirect targets across all conditions except MIN_Stat, LIV_Stat and RDM_Trans; a maximum of 8% of indirect targets can be explained by the currently known targets of direct Lrp targets (Table S4). Direct Lrp targets that are not currently identified as transcriptional regulators or regulators with incompletely documented regulons could account for why we are not able to explain more instances of indirect regulation, as could transcriptional units regulated by aspects of cellular state that are themselves Lrp-dependent. Several key transcription factors that are direct Lrp targets are responsible for explaining the identified indirect Lrp targets across conditions: Nac, GadW, PurR, LeuO, ArgR, QseB, CysB, NagC, SlyA, SoxS, and LrhA. Several of these transcription factors have also previously been identified as Lrp targets [28].

Investigating at a local as opposed to global scale provides several informative examples. At LIV_Log, LIV_Trans and RDM_Log, the dual regulator LrhA is a direct Lrp-activated target gene (Fig 3A). LrhA activates *fimE* and represses *flhC* and *flhD* (Fig 3B). At LIV_Log, *fimE* is indirectly activated; at LIV_Trans, *flhC* is indirectly repressed, and at RDM_Log,both *flhC and flhD* are indirectly repressed (Fig 3A). While this pattern does not show activity at every LrhA target in each condition, overall it suggests that indirect regulation of *fimE* and *flhCD* by Lrp may be explained in some cases by direct LrhA activation by Lrp. All three target genes are also known to be regulated by other transcription factors, potentially explaining the incomplete activity from LrhA. Similarly, at MIN_Trans, the transcriptional regulator CysB is a direct Lrp-repressed target gene (Fig 3C). CysB is known to activate *tcyP* and *cysI*, among other genes (Fig 3D). Both *tcyP* and *cysI* were categorized as indirect Lrp-repressed targets, supporting the fact that Lrp repression of *cysB* is what leads to repression of *tcyP* and c*ysI*.

**Figure 3:**
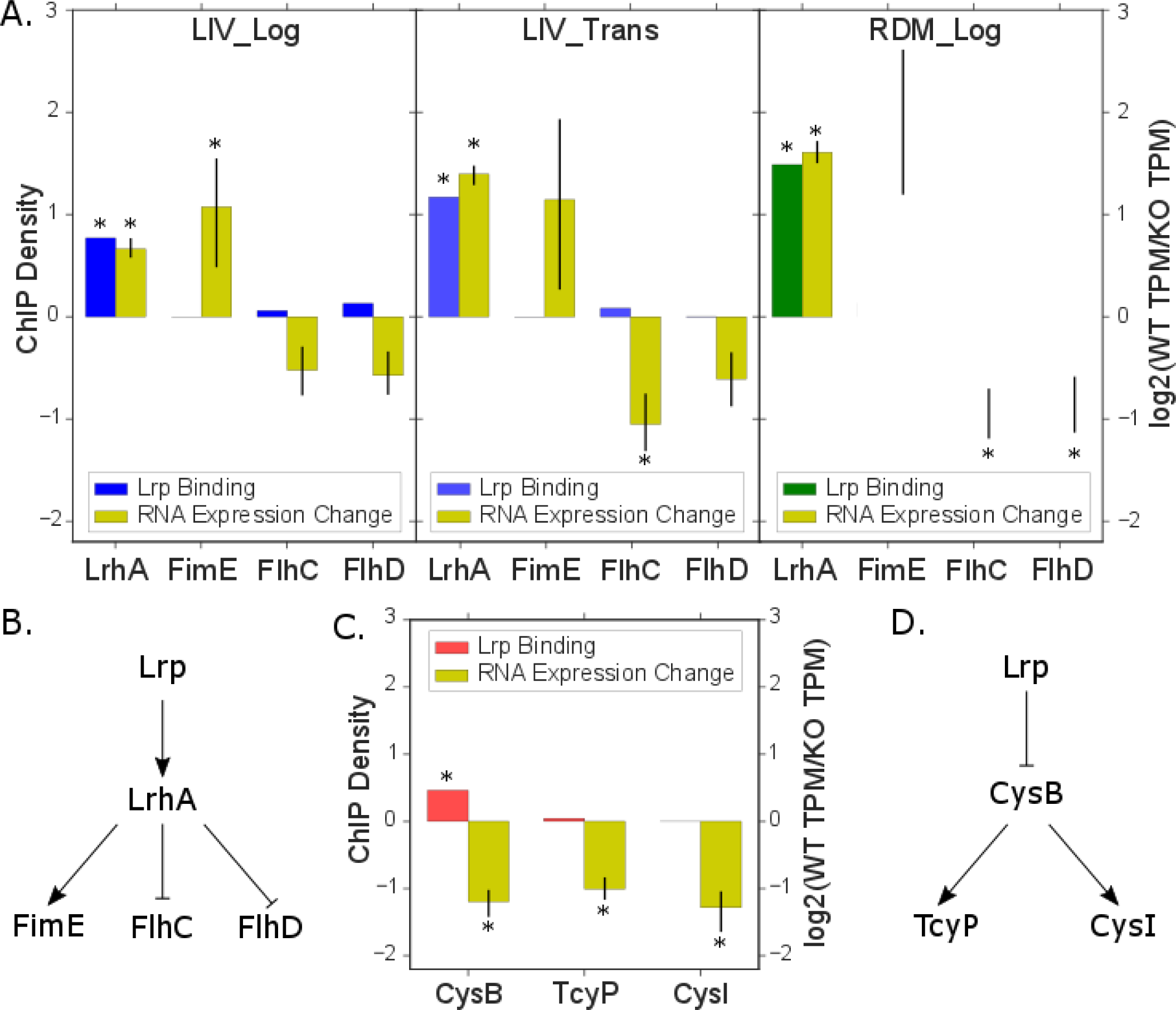
*Known targets of direct Lrp targets explain the mechanism of indirect Lrp regulation at some genes*. **A.** ChIP density and RNA-seq expression change (log2(WT/KO)) for direct Lrp target LrhA and its known target genes, FimE, FlhC and FlhD [18]. Error bars for the RNA-seq data indicate a percentile based 95% confidence interval from 100 bootstrap replicates of TPM estimates. Stars indicate a significant difference in RNA abundance between WT and *lrp::kanR* strains (Wald Test qvalue of < 0.05 and a genotype log fold change coefficient magnitude of > 0.5; see Methods for details). **B.** Proposed model of Lrp/LrhA mediated regulation of LrhA targets. **C.** ChIP density and RNA-seq expression change (log2(WT/KO)) for direct Lrp target CysB and some of its known target genes, TcyP and CysI [18], as in **A**. **D.** Proposed model of Lrp/CysB mediated regulation of CysB targets.

### Direct and indirect Lrp targets have both shared and unique GO-term classifications

After grouping direct and indirect targets, we used iPAGE [29] to search for enrichment of gene ontology (GO) terms that share mutual information with our categorization scheme. We observe the general trend that pathways involved in direct synthesis or acquisition of nutrients (e.g. amino acid transport and L-serine biosynthetic processes) tend to be direct targets or NAP-type targets, whereas those involved in regulation of cellular behavior and foraging strategies (e.g. flagellum and motility) tend to be indirect targets, particularly under the richer media conditions LIV_Log and RDM_Log (Fig 4A, Fig S2A). Interestingly, in testing for enrichment among the large block of indirect targets at RDM_Stat, we observe that it is depleted for flagellum-related genes. Under minimal conditions, indirect targets overlap with some of the transport pathways otherwise mainly observed to be enriched among direct targets.

**Figure 4:**
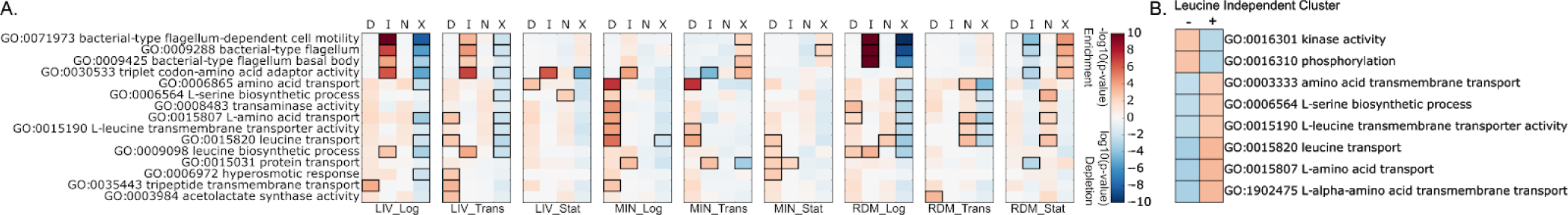
*Enriched GO-terms differ for direct and indirect Lrp targets*. **A.** A subset of GO-terms enriched or depleted within various conditions and groups of targets are listed to the left. Abbreviations are as follows: D - direct targets, I - indirect targets, N-NAP-type targets, X - no Lrp link genes. **B.** GO-terms enriched or depleted in the leucine independent cluster seen in Fig 2C. + indicates genes in the cluster and - indicates genes outside of the cluster. Boxes around a specific GO-term/condition/target group indicates a significant enrichment or depletion as indicated by a hypergeometric test (p-value < 0.01). Color inside the box specifies the magnitude of enrichment (red) or depletion (blue) as indicated by the color bar.

We also see overlapping enriched GO-terms at direct and NAP-type targets, suggesting that Lrp may preemptively bind some target genes before conditions occur at which regulatory action is required (discussed in more detail below). A particularly clear example of such poised regulation comes in identifying significant GO-terms among the genes that fall into the leucine-independent cluster from the categorization heat map; this cluster includes genes that are bound under almost all conditions, but only become indirect or direct targets during certain conditions. Strikingly, enriched GO-terms include leucine transport, serine biosynthetic processes and general amino acid transport (Fig 4B). This indicates that regardless of the level of leucine, Lrp’s traditionally recognized small molecular partner, Lrp remains bound to and poised to regulate critical genes if conditions change. Furthermore, the key signal causing a transition between NAP-type activity and direct transcriptional regulation is unlikely to be leucine levels themselves, as NAP to direct changes occur at certain genes across the time course of growth under Minimal conditions (see Fig 2C). Overall, the dynamic nature of what constitutes a Lrp-regulated target under different media conditions and points of growth demonstrates the complexity of the Lrp regulon.

In order to illuminate what distinguishes the various effects of Lrp binding on gene regulation, we tested whether splitting direct and indirect targets into sub-classes that are activated or repressed by Lrp revealed a different pattern of GO terms. From this analysis, it is evident that the flagellar genes are enriched among indirect Lrp-repressed targets specifically (Fig S2B). In addition, many of the genes involved in transport processes appear to be direct Lrp-repressed target genes, whereas the genes involved in biosynthetic processes are often direct Lrp-activated target genes. Interestingly, at MIN_Trans, a condition in which we see a spike of indirect targets, there is a specific class of GO-terms which are enriched for either indirect Lrp-activated (e.g. ferrous iron binding and N-terminal protein acetylation) or indirect Lrp-repressed targets (e.g. NAD binding and histidine biosynthetic processes). Those GO-terms do not have enrichment among indirect targets at RDM_Stat, reinforcing the notion that Lrp acts on unique sub-clusters of its targets under different conditions (as seen in Fig 2C above).

### Lrp is poised at many targets to enable combinatorial regulation

Upon filtering and categorizing genes, we noticed immediately that many genes shift between being a NAP-type target and a direct Lrp-activated or repressed target under different conditions (see Fig 2C above). In fact, 91% of direct Lrp targets are NAP-type targets in at least one condition, and thus have Lrp bound to their promoter even though it has no impact on transcription. For example, the MIN_Trans cluster consists of genes that are bound by Lrp in all Minimal media timepoints, but only show Lrp-dependent changes in transcript level during the Trans timepoint. This suggests that Lrp binds some promoters in a poised position under a broad range of conditions, but only regulates when certain additional criteria are met, perhaps by coordinating with a second regulatory factor to enable combinatorial logic. Among genes that undergo a transition between being a NAP-type target and a direct target, 38.8% become activated, 47.6% become repressed and 13.6% become both activated and repressed in different conditions.

For example, *potF*, a component of the putrescine ABC transporter [30], shows Lrp binding in its promoter region under all nine conditions measured in our data, but is only activated by Lrp during MIN, LIV_Stat, RDM_Log and RDM_Stat conditions (Fig 5A). In contrast with the variable Lrp-dependent RNA expression levels, Lrp binding at *potF* is very similar across conditions, spanning a similar length of DNA, and showing maximal signal at the same point. *potF* was previously identified as a Lrp regulated target which is repressed by Lrp alone, and activated when leucine is present [15]. However, those experiments employed glucose rather than glycerol as a carbon source, and monitored response to the addition of 10 mM leucine alone versus 0.2% (w/v) isoleucine, valine and leucine (equivalent to 15.25 mM leucine) which could explain the differences in observations of Lrp’s regulatory action at *potF*.

**Figure 5:**
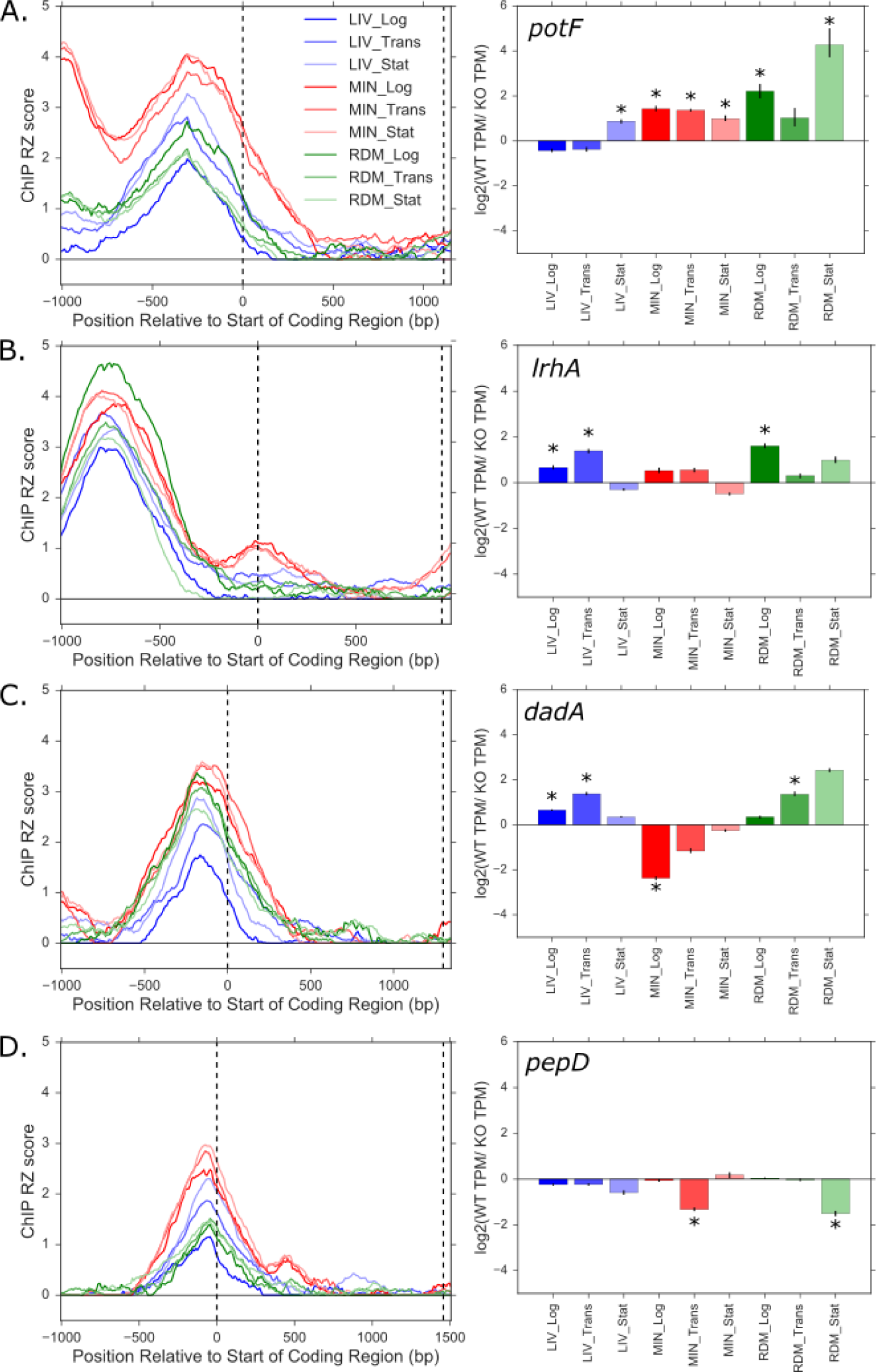
*Lrp sits at genes in poised position in preparation for regulatory activity*. ChIP robust Z-score (left) and RNA-seq expression change (log2(WT/KO); right) for four Lrp targets. *potF* (**A**) and *dadA* (**C**) are previously known targets, and *IrhA* (**B**) and *pepD* (**D**) are novel targets. Dashed vertical lines on the ChIP robust Z-score graph mark the start and end of the gene coding region. Error bars for the RNA-seq data indicate a percentile based 95% confidence interval from 100 bootstrap replicates of TPM estimates. Stars indicate a significant difference in RNA abundance between WT and *lrp::kanR* strains (Wald Test qvalue of < 0.05 and a genotype log fold change coefficient magnitude of > 0.5; see Methods for details).

*IrhA*, a transcriptional regulator involved in fimbriae synthesis [31], also has Lrp binding signal under all conditions. Interestingly, it is activated only at the high-leucine conditions LIV_Log, LIV_Trans and RDM_Log (Fig 5B), a different pattern from many other activated genes, which generally are activated in later time points in the growth curve or under MIN conditions. Again, the Lrp binding signal at *lrhA* is very similar across conditions, with only slight variation in the signal magnitude, in contrast to the sharp differences in the Lrp-dependent RNA expression changes. Thus, the changes in regulatory activity cannot be due to changes in the location of Lrp binding.

*dadA*, which encodes a critical enzyme in amino acid degradation [32], is one of the interesting class of examples that we see transition from a NAP-type target to being a direct Lrp-activated or repressed target in different conditions. *dadA* expression is strongly Lrp-repressed at MIN_Log, whereas it is activated during LIV_Log, LIV_Trans or RDM_Trans (Fig 5C). Lrp is known to repress *dadA* in the absence of leucine [33], a fact strongly supported by our data in which we see Lrp-mediated repression in minimal media and alleviation of repression during growth with higher levels of leucine. This variability in regulatory effect is in sharp contrast to the almost identical Lrp binding signal present in all nine conditions. Another gene which transitions between being a NAP-type target and being a direct Lrp-repressed target while having a similar Lrp binding profile is *pepD*, which is repressed at MIN_Trans and RDM_Stat (Fig 5D). *pepD* encodes Peptidase D, which cleaves a variety of dipeptides [34]. It is important to note that the location of the Lrp binding peak relative to the transcription start site does not systematically affect the direction of Lrp regulation (for example, compare *ilvI* and *potF*).

### Lrp connects with other regulatory factors

The phenomenon outlined above -- of Lrp frequently binding to a promoter under many conditions but only showing regulatory activity under a few -- suggests that other regulatory factors, such as σ factors or transcription factors, may be important in triggering an activating or repressive effect secondary to Lrp binding. If a σ factor and Lrp co-regulate some set of targets, we expect to see enrichment for direct targets relative to NAP-type targets within the σ factor’s regulon, especially at conditions when the σ factor is most active. To establish relative σ factor activity, we determined the average expression of all known σ factor target genes at each of our nine experimental conditions (Fig 6A) [18]. One caveat of our analysis is that some data is missing since we do not classify all genes in relation to Lrp, as outlined above, and, likewise, it is not known by which σ factor all genes that are classified are regulated. Subject to these constraints, our analysis in this section included 1534 genes. In addition, in some cases, overlap between other factors and Lrp may not indicate a direct interaction but may indicate that the other factor and Lrp have independent roles or functions at shared targets, here termed convergent regulation. However, if Lrp does interact directly with certain σ factors to activate target genes at specific conditions, there are a few possible explanations for why the NAP-type to direct target transition occurs at those points: 1) the transition only occurs when the genes’ controlling σ factor is active; 2) the nature or extent of Lrp binding itself changes at that condition; or 3) an accessory factor needed for Lrp-σ factor interaction is only present at that condition.

**Figure 6:**
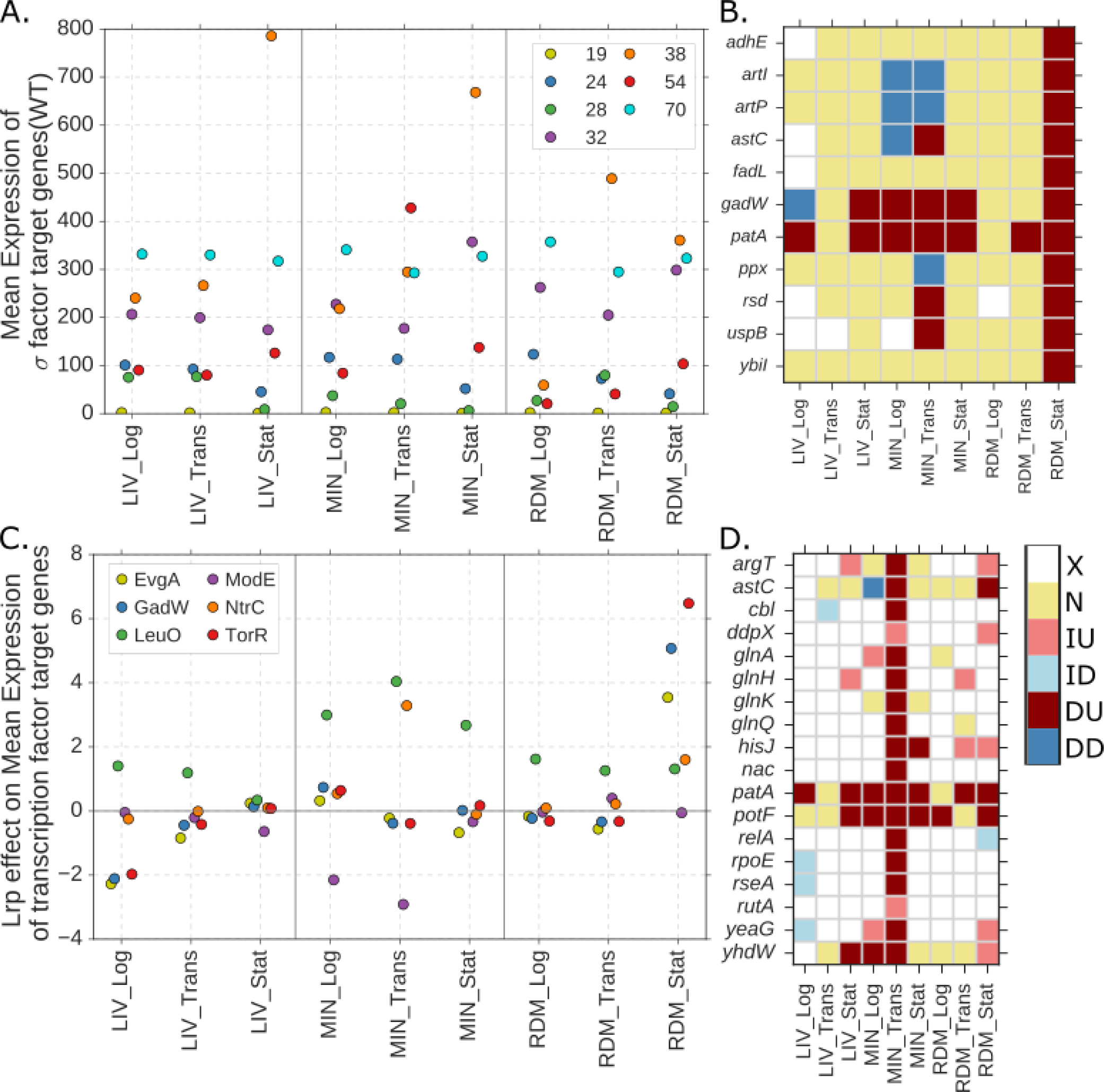
*Lrp interacts with other regulatory factors to control some targets’ expression*. **A.** Average expression (TPM) of known targets of each σ factor in WT cells at each condition. **B.** Heatmap showing classification of a subset of σ^38^ targets which are direct Lrp-activated targets at RDM_Stat. Abbreviations on the color bar are as follows: DD - direct downregulated targets, DU - direct upregulated targets, ID - indirect downregulated targets, IU - indirect upregulated targets, N - NAP-type targets, X - no Lrp link. **C.** Average log2(WT/KO) expression ratio of known transcription factor targets for selected transcription factors at each condition. **D.** Heatmap showing classification of those NtrC targets which have an annotated transcription start site and thus are classified in our analysis. Abbreviations as for **B**.

We applied a permutation test to identify any σ factors with a significant enrichment of overlap between their targets and all direct Lrp targets or specifically direct Lrp-activated targets. All q-values and enrichment levels for the permutation test with all direct targets are listed in Table S5; results from the permutation test with only direct-activated targets are in Table S6 (r=10000 for both). Only two σ factors had significant overlaps: σ^54^ at MIN_Trans and σ^38^ at RDM_Stat. A role for σ^38^ at stationary phase is logical since it coordinates general stress responses in *E. coli [35]*. However, direct σ^38^/Lrp interaction is not likely since many of the Lrp-σ^38^ shared regulated genes at RDM_Stat are NAP-type Lrp targets in other conditions (MIN_Stat, LIV_Stat and RDM_Trans) when σ^38^ is more active (Fig 6A,B). Therefore, this overlap is likely a result of convergent regulation between Lrp, a “feast-famine” regulatory protein, and σ^38^, a stress-response σ factor.

In contrast with σ^38^, for σ^54^ we observe marked enrichment, especially of direct Lrp-activated targets, under the condition when σ^54^ is most active. Specifically, we document enrichment for direct Lrp targets with σ^54^(σ^N^) at MIN_Trans (1.8-fold enrichment, q-value: 0.092). At MIN_Trans, 39 % of Lrp binding sites overall are direct targets, whereas 68 % of σ^54^ targets with Lrp binding sites are direct targets. Furthermore, as we would expect for the case where Lrp acts as a co-activator for a given σ factor, there is enrichment specifically for direct Lrp-activated target genes among σ^54^ targets at MIN_Trans (3.0-fold enrichment, q-value: 0.022). Overall, 20% of Lrp binding sites are direct activated targets at MIN_Trans, whereas Lrp-bound targets in the σ^54^ regulon are direct Lrp-activated targets 59% of the time, a 3.0-fold increase. σ^54^ regulates many genes involved in nitrogen assimilation [36], and these results indicate that Lrp is likely involved in co-activating some σ^54^ dependent genes, in agreement with Lrp’s role in sensing and responding to nutrient levels. At MIN_Trans, Lrp actually also weakly represses σ^54^ itself directly; σ^54^ is not a direct or indirect target under any other conditions (Fig S3B).

Average expression of σ^54^ targets peaks at MIN_Trans (Fig 6A), in agreement with when we see overlap between its targets and direct Lrp-activated targets (12.5% of the direct Lrp-activated targets at MIN_Trans are known σ^54^ targets, and conversely 21% of the classified σ^54^ targets are direct Lrp-activated targets at MIN_Trans). Nine out of the thirteen overlapping target genes only become a direct Lrp-activated target at MIN_Trans. The remaining four genes (*astC, hisJ, potF, yhdW*) are affected at other conditions when there is a slight peak in σ^54^ activity, as measured by the overall expression of known target genes (Fig 6A), and could be subject to other regulatory control. For example, *astC* and *hisJ* are also regulated by ArgR in some conditions [37,38]. The fact that the shared regulated genes are only direct Lrp-activated targets when σ^54^ itself is most active supports the notion that σ^54^ may require Lrp binding to activate transcription of certain genes. At a molecular level, this suggests that while expression of σ^54^ itself during MIN_Trans does not require Lrp (and in fact, is slightly repressed by Lrp), its transcriptional activity is enhanced by the presence of Lrp (also see Fig S3A).

To investigate the possibility that Lrp binding itself changes to facilitate interaction with σ^54^, we visualized the Lrp-ChIP binding signal at shared direct Lrp/σ^54^ targets. Changes in Lrp binding, either complete reversals of binding between conditions or changes in peak length, are evident in the cases of some genes (*glnH, yeaG* and *yhdW*), while others, such as *ibpB* and *potF* have almost identical binding regardless of condition (see Fig 5A for *potF* Lrp-binding signal and Supplementary Data File 1); thus, it is unlikely that changes in Lrp binding itself are in general responsible for the regulatory interaction with σ^54^. Given that σ^54^ is known to require activating factors, it is likely that an accessory factor may facilitate Lrp/σ^54^ co-regulation.

To identify other candidates for co-regulators acting with Lrp, just as we tested for Lrp co-regulation with σ factors, we investigated whether Lrp has particular correlations with any of the other annotated transcription factors in *E. coli*. We compared the average expression of all annotated targets of individual transcription factors in WT and Lrp KO conditions to identify those transcription factor regulons that show Lrp-dependent changes. Several transcription factors were identified as significant based on a permutation test (r=1000): ArcA, CsgD, EvgA, FlhDC, GadW, LeuO, ModE, NtrC and TorR. We then applied the additional threshold of requiring an average four-fold or greater change in expression of target genes dependent on Lrp status (WT vs. KO) to identify the most biologically relevant interactions (Fig 6C); the transcription factors ArcA, CsgD and FlhDC did not pass this filter and were eliminated from further analysis. EvgA, LeuO, ModE and TorR all likely represent convergent regulation due to the existence of no or limited overlap between transcription factor targets and direct Lrp targets. The transcription factor GadW’s Lrp dependency at LIV_Log is likely due to it itself being a direct Lrp-repressed target (Fig S3C) -- in investigating the classification of genes in its regulon, we see over half are indirect Lrp-repressed genes at LIV_Log. Like the examples of LrhA and CysB shown above, we again see that direct regulation of a transcription factor by Lrp can mediate changes in many other genes.

The transcription factor NtrC is a notable exception to the above trends, as 33% of all its targets are also direct Lrp-activated targets (Fig 6D). This number is an underestimate since it only accounts for the genes classified in our scheme (namely those with annotated promoters); if we expand our classification to include the genes that comprise the transcription units of those classified genes, 74% of NtrC targets are also direct Lrp-activated targets. Two indirect Lrp-activated transcription units comprise the remainder of the NtrC regulon. NtrC is one of the transcription factors which can serve as an activator of σ^54^, so the intersection between Lrp, NtrC and σ^54^ is interesting to consider. Activators of σ^54^ often bind at a distance from the promoter and so require significant DNA bending to come in physical contact with σ^54^ [39]. IHF is a DNA bending protein known to facilitate DNA bending at some target genes, but our data indicates that Lrp may also have a role in DNA bending, and thus activation of NtrC/σ transcribed genes (see Discussion). Thus, while many instances of Lrp regulation appear to require co-regulation with as yet unidentified regulatory factors, we are able to identify some likely possible mechanisms.

### Lrp binding sites have a condition- and time-specific motif preference

While not as invariant as motifs for other *E. coli* transcription factors, a 15 base-pair motif comprising terminal inverted repeats and an AT-rich center was previously identified for Lrp [15,40]. We wanted to determine if a similar motif is apparent in our data, and how well Lrp binding is predicted by the presence of Lrp motifs. We used a logistic regression model to classify 500 bp windows of the genome as either containing a Lrp peak or not, using as predictors the presence of previously documented Lrp motifs and the AT content (given the AT richness of the Lrp motif itself). Starting with a minimal model containing only an intercept term, we created more complex models by adding a single predictor at a time and scoring each new model with the Bayesian Information Criterion (BIC) as displayed in Fig 7A; *n.b.* a lower BIC indicates a more parsimonious model. A minimal model was chosen by adding to the new model the predictor with the largest decrease in BIC from the intercept-only model and iterating this process until the change in BIC switched sign (indicating that additional terms were no longer informative). A similar analysis was done in which we started with a full model containing all of the predictors and removed the predictor with the largest increase in BIC until the change in BIC switched sign (Fig S4). In both cases we arrived at the same set of minimal models for each condition. Intriguingly, among the minimal models for each condition, we see a shift between a non-specific preference for AT-rich regions at Log points and specific motif preference at later time points across all conditions (Fig 7A). In each condition, from early to late timepoints, there is a decrease in how much information is provided by the AT-content in terms of predicting Lrp binding. While their relative importance to the model shifts, the minimal variables needed to explain most of the data include a combination of AT-content and established Lrp motifs across all conditions. This suggests that Lrp binding is more non-specific in earlier phases of growth, and only gains specificity upon nutrient limitation and entrance into stationary phase, which also agrees with our observed increase in the number of peaks in later time points. Additionally, this pattern of specificity agrees with Lrp’s proposed position of importance as a regulator of the transition to stationary phase. However, since we see the same lack of specificity in LIV_Log and MIN_Log (two conditions with dramatically different leucine concentrations), we can conclude that leucine level alone is not sufficient to shift the binding specificity of Lrp, but rather, that other signals (such as, potentially, energy/carbon source availability) must also be integrated somehow into Lrp’s binding.

**Figure 7:**
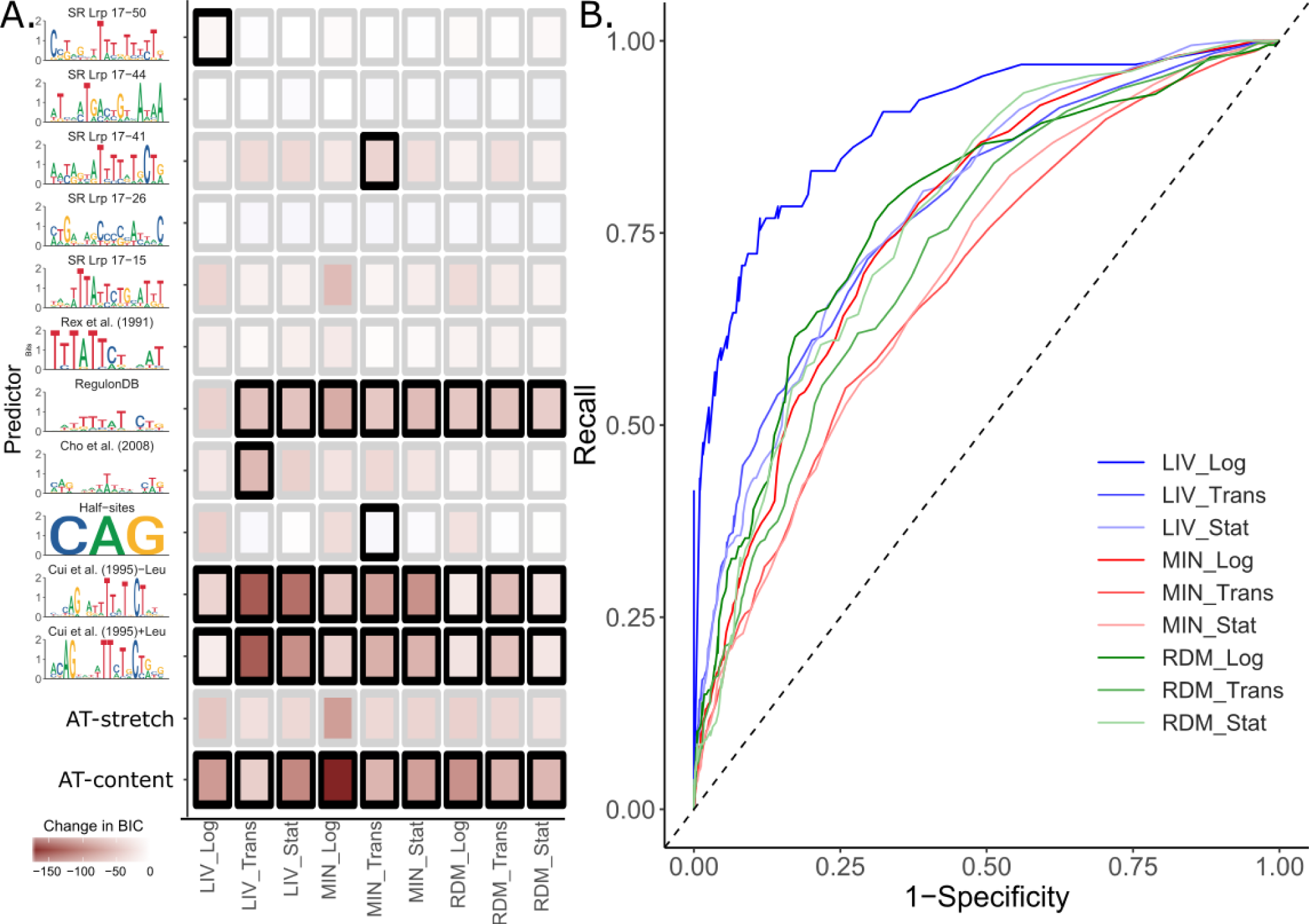
*Lrp exhibits condition-dependent sequence-preference*. **A.** Change in BIC for add-one-in logistic regression models. The y axis displays the Position Weight Matrix (PWM) used to create a particular feature. PWMs were obtained from the publication indicated above the PWM [15,40,71], RegulonDB [18] or, in the case of SR motifs, the SwissRegulon [72].Features were created from a given PWM by dividing the count of matches within a sequence (as obtained by FIMO [73] with p-value < 0.0001) by the length of the sequence. AT-stretch indicates the longest stretch of continuous As and Ts normalized by the length of the sequence. AT-content indicates the number of As and Ts normalized by the length of the sequence. Colors then indicate the change in BIC when a given term is added to a minimal model containing only an intercept term. Heavy boxes indicate a feature was included in the final model for that condition. For both this panel and panel **B**, the positive class of sequences was obtained by taking 500 bp around the center of each peak for each condition. The negative class of sequences was obtained by taking three times the number of equal-sized random sequences from the subset of the genome that was not in a peak for that condition. **B.** Receiver Operator Characteristic curves for each final model by condition. Curves were calculated at 0.01 increments from 0 to 1 for a predicted probability cutoff from the logistic regression. Full statistics including five-fold cross-validation are included in Table S7.

The performance of the derived models is relatively good; the receiver operator curves, which show the recall for every potential false positive rate, trend toward the upper left corner where a perfect model would be (Fig 7B; quantified by area under the curve, ROC-AUC, in Table S7). In addition, the Matthews correlation coefficient (MCC), a combined measure of precision and recall which has potential values from 0 to 1, ranges from 0.25 to 0.60 (Table S7). These performance metrics were robust to withholding of shuffled subsets of the data, as indicated by minimum and maximum values found in five-fold cross-validation (values in parenthesis in Table S7). Overall the specificity of these models is much better than their sensitivity, indicating that they perform well in rejecting locations where Lrp does not bind. However, there is still substantial room for improvement in calling Lrp bound sequences. Interestingly, the sensitivity drops in the conditions where specific sequence motifs are more informative. It is likely that we are missing additional features that would improve the sensitivity in these conditions; however, efforts to discover additional sequence determinants of Lrp binding were unsuccessful. This could simply indicate that sequence independent mechanisms, such as the well-established observation of Lrp cooperativity in binding [41], or recruitment of Lrp by binding of additional factors, could play a role in determining Lrp binding locations.

## Discussion

### Lrp regulates hundreds of genes in distinct categories by direct and indirect mechanisms

By investigating Lrp activity under several media conditions and timepoints, and integrating binding data with changes in RNA expression, we are able to present an enhanced view of the Lrp regulon. Our use of a high-quality antibody against native Lrp removes any possibility of epitope tagging hindering native behavior in our experiments, and the use of modern sequencing-based methods provides us with a high resolution snapshot of both Lrp’s binding and regulatory activity. We document hundreds of novel targets, and note the especially important effect of indirect regulation at MIN_Trans and RDM_Stat. The differences between direct and indirect targets are borne out by the GO-term analysis in which we see a shift between GO-terms at direct targets (more transport and biosynthesis related genes) and those at indirect targets (flagellum associated genes among others). This could point to organization at a temporal level; the genes needing most urgent regulation (such as those involved directly in importing or generating needed nutrients) may be under direct Lrp control, while genes requiring less urgent modulation and instead governing foraging strategies may be indirectly regulated by Lrp.

In the most straight-forward transcriptional regulatory system, indirect targets should be traceable to a direct target. However, the complicated, interconnected nature of the regulatory system of *E. coli* may explain why we are unable to find connections explaining all Lrp indirect targets. In some cases, there may be another layer of regulation before indirect Lrp targets are affected, or intracellular signaling pathways may be triggered, leading to broader downstream effects, such as changes in metabolite levels. The cases of CysB, LrhA and GadW cleanly illustrate how some indirect regulation is accomplished. Other cases of missed identification may also arise simply due to our incomplete knowledge of the regulons of all *E. coli* transcription factors.

### Primed Lrp binding argues for interaction with co-regulatory factors

From our experiments, we identify many points at which Lrp binds the regulatory region of a gene without producing an effect on transcription, and even points at which an apparently identical Lrp binding pattern has no effect on transcription in one condition, but has a substantial effect under another. Given that Lrp binding is enriched in regulatory regions relative to other locations in the genome, this argues against a purely DNA-organizing role for these NAP-type sites. If that was the case, we would expect Lrp binding sites (the majority of which are NAP-type sites in any condition) to be distributed more evenly across the genome. This poised regulation is also seen for some eukaryotic transcription factors such as the tumor suppressor p53 in binding to the *mdm2* gene [20]. Therefore, while Lrp itself is not conserved in eukaryotes, its ability to bind without regulating may have parallels to eukaryotic regulation, suggesting convergent evolution to a similar regulatory scheme. There are several possibilities for why Lrp may not have regulatory function in all cases where it binds, including 1) Lrp acts as a scaffold to interact directly with other proteins which are only present at certain conditions and modulate transcription, 2) Lrp wraps DNA in order to control DNA accessibility of other regulators, reminiscent of eukaryotic histone-like behavior, and/or 3) the presence of Lrp octamer or hexadecamer may control or influence the regulatory behavior of Lrp. We investigated the first possibility by analyzing if certain σ factors or transcription factors might be responsible for the condition-dependent regulation on a global scale. Although we do not see strong global evidence, gene-level studies have previously implicated Lrp in interacting with σ^38^ [42,43]. While many potential connections appear to be cases of convergent regulation, we identified a few specific cases where Lrp appears to play a direct role in modulating the effects of other regulators.

There are several data points that indicate direct interaction between Lrp and σ ^54^at MIN_Trans. First, σ^54^ is most active globally at MIN_Trans, in agreement with when we see many of the overlapping regulated genes transition from NAP-type to direct targets. As noted above, σ^54^ is unique among the *E. coli* σ factors in that it requires an activator, such as NtrC or PspF [44]. We also document enrichment for NtrC targets at MIN_Trans which argues for a role for Lrp in the nitrogen-limitation response. Known NtrC targets account for 33% of genes in the σ^54^ regulon, and almost all of those targets are in operons directly controlled by Lrp. Activators of σ^54^, such as NtrC, often bind to an upstream site and require precise looping of the DNA in order to bring the activator in contact with σ^54^; in previous studies, the bending has been documented as being intrinsic to the region or looping mediated by IHF [45]. In accordance with the possibility of intrinsic bending, the average AT content upstream of σ^54^ target genes is 70%, with the lowest being at 50% [36]. As previously reported and seen in our data, Lrp is known to bind AT-rich regions preferentially [46]. Lrp induces bending of 52° to 135° depending on the size of the binding sites [47]. Thus, we hypothesize that Lrp may play a role in bending DNA to coordinate NtrC-σ^54^ interaction at NtrC targets. This would agree with the connection between Lrp and nitrogen metabolism regulation seen previously in genome-wide studies [48]. Analogous interactions with other transcription or regulatory factors may explain other NAP-type/direct target transitions. For example, Lrp interaction with H-NS is important for regulating rRNA promoters [49], and Lrp competition with DNA adenine methyltransferase is critical in regulating expression of the *pap* operon, which produces pili [50]. In addition, non-protein small molecules like ppGpp are known to affect some Lrp-regulated target genes [51]. Further studies are needed to investigate Lrp’s interactions with other regulatory factors and the alternate mechanisms proposed above.

### Lrp binding activity is partially predicted by known sequence motifs

While we identify a preference for Lrp binding at several related motifs and AT-rich regions, there are still a significant subset of peaks that are not predicted by these models. Attempts to improve Lrp binding prediction from additional sequence determinants were not successful despite application of several state-of-the-art motif finders. As mentioned above, this could be due to Lrp binding initially at a sequence-specific location, and subsequent Lrp molecules binding due to cooperativity and the high local concentration of Lrp molecules provided by Lrp’s oligomeric nature. Alternatively, Lrp itself may be recruited by other proteins. Due to Lrp’s relatively high non-specific DNA binding affinity, especially under rich conditions [14], it is reasonable to find that not all of its binding locations can be predicted based on sequence alone. It is again important to note that the switch in DNA-binding specificity occurs regardless of the levels of leucine, suggesting that other small molecule regulators [16] or potentially post-translational modifications [52,53] may play a role in Lrp regulatory activity. Additionally, despite extensive effort, we were unable to identify any sequence determinants capable of reliably explaining Lrp regulatory activity, either through predicting transitions from poised to active regulation, or distinguishing Lrp activation from Lrp repression. Possible mechanisms for this behavior include interactions with condition-specific factors that bind near the multifunctional Lrp sites (many potential partners have likely not yet been characterized), condition-dependent DNA looping triggered by the binding of Lrp to nearby sites or by octamer-hexadecamer transitions, or post-translational modifications to Lrp itself. Dissecting the detailed molecular mechanisms underlying the binding and regulatory landscape that we have revealed here will be a fruitful area for future research.

## Methods

### Strains and media

The WT strain used in this study was *E. coli* K-12 MG1655 (ATCC 47076). The Lrp deletion strain was constructed by homologous recombination resulting in the insertion of kanamycin resistance cassette [54]. Primers used for strain construction and validation are listed in Table S8. The *lrp::kanR* strain was validated by sizing of the P965/P1568/P1569 products and Sanger sequencing.

All routine cell growth during cloning was done in LB medium (10 g/liter tryptone, 5 g/liter yeast extract, 5 g/liter NaCl) or on LB plates (LB medium plus 15 g/liter Bacto agar) supplemented with 50 μg/mL kanamycin or 100 μg/mL ampicillin (both from US Biological; Salem, MA) as required. For the ChIP-seq and RNA samples, a single colony of wild type *E. coli* or the *lrp::kanR* strain was inoculated into MOPS media (Teknova; Hollister, CA) with 0.04 % glucose [55] and grown overnight. The cells were then back-diluted to OD600=0.003 in 100 mL of the appropriate target media. Experiments were performed in MOPS with 0.2 % glycerol (the MIN media condition), MOPS with 0.04 % glycerol and 0.2 % (weight/volume) each leucine (Amresco; Solon, OH), isoleucine (Alfa-Aesar; Haverhill, MA) and valine (Amresco; Solon, OH; the LIV condition), or MOPS plus 0.4% glycerol, ACGU and EZ supplements (Teknova; Hollister, CA; the RDM condition). Media conditions are summarized in Table S9.

The cells were grown at 37°C with shaking (200 rpm) until the OD600 was between 0.15 and 0.25 (for log phase samples), between 1.8 and 2.2 (for transition point in MIN or LIV media), between 2.3 and 2.7 (for transition point in RDM), or 12 hours past the log point (for stationary phase samples). The OD600 range for transition point harvest was determined by monitoring the growth of cells grown in conditions identical to the experiment and selecting the point in the OD600 range during which exponential growth becomes non-linear when visualized on a log scale.

### ChIP-seq

At the appropriate time, either WT or *lrp::kanR* cells were cross-linked by adding formaldehyde (37 % Sigma-Aldrich; St. Louis, MO) to a final concentration of 1 % (vol/vol) and incubated with shaking for 15 minutes at room temperature. Formaldehyde cross-linking was quenched by addition of Tris (pH 8) to a final concentration of 280 mM and incubation with shaking at room temperature for 10 minutes. The culture was then immediately centrifuged for 5 minutes at 5500 xg at 4 ° C. The pellet was washed twice with 30 mL ice cold TBS (50 mM Tris, 150 mM NaCl, pH 7.5) before being resuspended in 1 mL TBS. Following a 3 minute centrifugation at 10,000xg at 4°C and removal of the supernatant, the pellet was flash-frozen in a dry ice/ethanol bath and then stored at −80°C. Two biological replicates, grown on different days, were prepared for each condition.

The cell pellet was resuspended in lysis buffer (PBS, 0.1 % Tween 20, 1 mM EDTA, 1x Complete Mini EDTA-free Protease Inhibitors (Roche; Basel, Switzerland), 0.6 mg lysozyme (Amresco; Solon, OH)), vortexed for 3 seconds, and incubated at 37°C for 30 minutes. The sample was then sonicated in 3 bursts of 10 seconds each at 25 % power (Branson Digital Sonifier). Cellular debris was removed by centrifugation at 16,000xg for 10 minutes at 4°C. As an input sample, 50 uL of the supernatant was removed and mixed with EDTA to 8.6 mM and 235 uL Elution Buffer (50 mM Tris (pH 8), 10 mM EDTA, 1 % SDS (vol/vol)). The remainder of the lysate was added to 50 uL pre-washed SureBeads Protein G magnetic beads (Bio-Rad; Hercules, CA) and rocked for 1 hour at room temperature for pre-clearing. A separate aliquot of 100 uL of pre-washed SureBeads Protein G magnetic beads was incubated with 10 ug Lrp monoclonal antibody (Neoclone; Madison, WI) for 10 minutes at room temperature with rocking and then washed thrice with PBS/0.1 % Tween-20 before the pre-cleared supernatant was added. The bead/lysate mixture was again incubated with rocking for 1 hour at room temperature. The beads were then washed thrice with PBS/0.1 % Tween-20. To elute the cross-linked Lrp/DNA complexes, the beads were resuspended in 285 uL of Elution Buffer and incubated at 65°C for 20 min, vortexing every 5 minutes. The resulting eluate was incubated overnight at 65°C to reverse the cross-links.

The sample was treated with 0.05 mg RNase A (Thermo Fisher; Waltham, MA) for 2 hours at 37°C, then 0.2 mg Proteinase K (Thermo Fisher; Waltham, MA) for 2 hours at 50°C before the DNA was isolated by phenol-chloroform extraction and ethanol precipitation. The samples were quantified (QuantiFluor dsDNA Kit, Promega; Madison, WI) and prepared for sequencing using the NEBNext Ultra DNA Library Prep Kit for Illumina (NEB; Ipswich, MA). The library was checked for quality by 2 % agarose gel electrophoresis using GelRed stain (Biotium; Fremont, CA). Samples were pooled and the sequencing performed on an Illumina NextSeq −500, with 38×37 bp paired end reads. We obtained at least three million reads that passed all filters and aligned properly to the genome per biological replicate with an average of nine million reads per replicate (Table S10). Input samples were treated identically to the ChIP extracted samples beginning at the RNase A treatment.

### RNA-seq

For RNA-seq samples in both WT and *lrp::kanR* cells, 2.5 ml of culture was removed when cells had reached the appropriate OD and mixed with 5 mL Qiagen RNAProtect Bacteria Reagent (Qiagen; Hilden, Germany), vortexed, incubated 5 minutes at room temperature, and then centrifuged for 10 minutes at 5,000 xg in a fixed angle rotor at 4°C. The supernatant was removed and the pellet was flash-frozen in a dry ice/ethanol bath before being stored at −80°C.The pellet was resuspended in TE and treated with 177 kilounits Ready-Lyse Lysozyme Solution (Epicentre; Madison, WI) and 0.2 mg Proteinase K (Thermo Fisher; Waltham, MA) for ten minutes at room temperature, vortexing every two minutes. The RNA was purified using the Zymo RNA Clean and Concentrator kit (Zymo; Irvine, CA), treated with 5 units Baseline Zero DNase (Epicentre; Madison, WI), in the presence of RNase Inhibitor (NEB; Ipswich, MA), for 30 minutes at 37°C, and then again purified with the Zymo RNA Clean and Concentrator kit. RNA quality was assessed by electrophoresis in a denaturing agarose-guanidinium gel [56]. rRNA depletion was performed using the Ribo-Zero rRNA Removal Kit for Bacteria (Illumina; San Diego, CA), halving all reagent and input quantities but otherwise following the manufacturer’s instructions. cDNA synthesis and sequencing library preparation were performed following the NEBNext Ultra Directional RNA Library Prep Kit (NEB; Ipswich, MA). The library was checked for quality by 2% agarose gel electrophoresis using GelRed stain (Biotium; Fremont, CA).Samples were pooled and the sequencing performed on a NextSeq −500 at the University of Michigan’s DNA Sequencing Core Facility.

### Preprocessing of ChIP-seq data

Sequencing adapters were removed from all sequences using CutAdapt version 1.8.1 [57] with parameters -a AGATCGGAAGAGC -A AGATCGGAAGAGC -n 3 -m 20 --mask-adapter --match-read-wildcards. Low quality reads were trimmed with Trimmomatic version 0.32 [58] using the parameters TRAILING:3 SLIDINGWINDOW:4:15 MINLEN:20. The quality of the raw and preprocessed fastq files was assessed using FastQC version 0.10.1 [59] and MultiQC version 1.2 [60]. The number of raw and surviving reads for each sample are described in Table S10.

### Alignment of ChIP-seq data

All samples were aligned to the MG1655 U00096.2 genome modified to match the insertions and deletions for the ATCC 47076 variant of E. coli MG1655 as reported by [61]. Alignments were performed using bowtie version 2.1.0 [62] and arguments: -X 2000 -q --end-to-end --very-sensitive -p 5 --phred33 --dovetail in order to maximize the sensitivity of the alignment. Final alignment rates for each sample are described in Table S10.

### Calculation of ChIP-seq summary signal

The coverage *c* of paired-end reads at every tenth base pair *n* across the genome, was calculated from the alignments for the ChIP-extracted and input tracks for each sample and replicate separately using custom python scripts. The raw coverage for extracted and input signals was then scaled using the median coverage across the genome for each individual track. The raw enrichment (RE) was calculated using the log2 ratio of scaled extracted to input coverage separately for each pair of extracted and input samples as shown below:

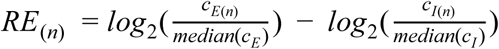

Where *E* and *I* denote the extracted and input samples, respectively. Since the Lrp WT and *lrp::kanR* samples are not paired, each combination of raw Lrp enrichment for a Lrp WT replicate and a *lrp::kanR* replicate was used to generate a raw subtracted enrichment signal (RSE) for the Lrp WT - *lrp::kanR* signal, resulting in four possible subtracted replicates for each condition and time point. The *lrp::kanR signal* was subtracted only if it was positive.

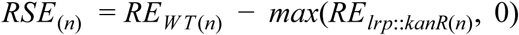

For each replicate pairing, the raw subtracted Lrp enrichment signals were converted to robust Z-score estimates (RZ) using the following formula:

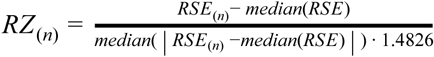

The RZ replicates were then averaged to generate the final occupancy signal for downstream analysis. Reproducibility of both the RE and RSE for each replicate can be seen in Fig S5A.

### Determination of high-confidence Lrp binding sites

In order to determine regions of high-confidence Lrp binding we required three criteria for Lrp enrichment to be satisfied: 1. The enrichment must be technically reproducible. 2. The enrichment must be above the input background. 3. The enrichment must be biologically reproducible. The following paragraphs detail how each of these criteria were determined.

#### Assessment of technical reproducibility of Lrp enrichment

To assess the technical reproducibility of the Lrp enrichment, we used custom python scripts to sample with replacement from the aligned reads separately for each paired extracted and input sample. The RSE for each of the four possible subtracted Lrp WT vs. *lrp::kanR* replicates was calculated, as described above, for each of 1000 bootstrap replicates. To test for technically reproducible enrichment, we considered a null hypothesis that the RSE is normally distributed centered at 0. A Z-score for each location *n* was then determined as follows

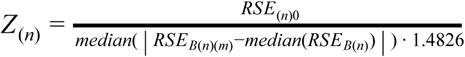

Where *RSE*_0_ is the unsampled dataset and *RSE*_*B*_ represents the bootstrap replicates for which *m* = 1: 1000. The resulting Z score was converted to a p-value using a one-sided Z test through the scipy.stats normal cumulative distribution function [63]. These p-values were FDR corrected using the procedure described by Benjamini and Hochberg [64]. A region was considered to be technically reproducible if its q-value was less than 0.001.

#### Assessment of Lrp-specific enrichment

To assess enrichment of ChIP signal above the input background and to differentiate from off-target antibody enrichments seen in pulldowns using the *lrp::kanR* strain, an *RZ* score (see above) was calculated for each combination of WT-*lrp::kanR* replicates, yielding positive signal only when the WT pulldown value was substantially above that of the *lrp::kanR signal*. We then tested for enrichment of the RZ score above the median signal for that track through the use of a one-sided Z-test using scipy.stats normal cumulative distribution function and FDR correction of the resulting p-value to a q-value. To be considered enriched above background, a region was required to have an enrichment qvalue less than 0.001.

#### Assessment of biological reproducibility

To assess the biological reproducibility of each region *n,* the irreducible discovery rate [65] was calculated for each data point between the RZ signals for each of the four Lrp WT - *lrp::kanR* combinations for each condition and timepoint. Starting parameters for the IDR calculation for each condition included °= 0.0, σ = 1.4826, ρ = 0.1 and an associated weight based on the estimated number of bound Lrp octamers for each nutrient condition *x* as estimated in [14]:

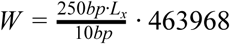

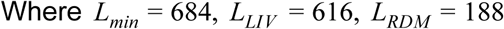

A region was considered to be biologically reproducible if the FDR-corrected IDR q-value was less than 0.01 for both combinations of RSE replicates.

### Combining enrichment and reproducibility into final peaks

Final peaks were determined if a region *n* passed the biological reproducibility filter and at least one of the four subtracted replicate combinations passed both the technical and enrichment filters. Adjacent passing regions were consolidated into one region if they were within 30 base pairs. The applied cutoffs and other thresholds were confirmed to be reasonable through manual inspection of called peaks and candidate peaks that narrowly missed one or more cutoffs. An example peak in comparison to a non-Lrp-specific peak can be seen in Fig S5B,C.

### Preprocessing of RNA-Seq data

Similar to the ChIP-Seq reads, sequencing adapters were removed from all sequences using CutAdapt version 1.8.1 [57] with parameters --quality-base=33 -a AGATCGGAAGAGC -A AGATCGGAAGAGC -n 3 -m 20 --mask-adapter --match-read-wildcards. Low quality reads were trimmed with Trimmomatic version 0.32 [58] using the parameters LEADING:3 TRAILING:3 SLIDINGWINDOW:4:15 MINLEN:20. The quality of the raw and preprocessed fastq files was assessed using FastQC version 0.10.1 [59] and MultiQC version 1.2 [60]. The number of raw and surviving reads for each sample are described in Table S11.

### Filtering highly abundant RNAs from analysis

In some, but not all, of our samples as much as 70% of our RNA-seq reads were ribosomal reads or the highly abundant RNA products from *ssrA* and *ssrS*. (Table S11). To filter highly abundant RNA reads and thus avoid having variations in ribosome depletion efficiency interfere with proper normalization, all RNA-seq reads were aligned using bowtie2 version 2.1.0 [62] to the same ATCC 47076-modified version of the U00096.2 genome used for the ChIP-Seq data. The following parameters were used for bowtie2: -q --end-to-end --very-sensitive -p 5 --phred33 --dovetail. The subsequent alignments were parsed for reads that overlapped with ribosomal reads in a strand specific manner using custom python scripts. New fastq files were written that only included reads that did not overlap ribosomal reads, and these files were used for downstream gene expression analyses. In all replicates at least two million reads survived this final filter with the smallest size replicate containing 2.6 million reads after filtering (Table S11).

### Gene-centric quantification of RNA-Seq Data

Gene-centric quantification of RNA expression for all samples was performed using kallisto version 0.43.0 [66] with the arguments: quant -t 4 -b 100 --rf-stranded. The appropriate transcriptome file needed for alignment was created through converting the GeneProductSet dataset from RegulonDB version 9.4 [18] to the appropriate ATCC 47076 coordinates and input file format for kallisto using custom python scripts.

### Determination of Lrp-dependent changes in Transcription

To determine Lrp-dependent changes in transcription, we used kallisto’s companion post-processing data analysis software, sleuth [67] to model the transcript abundance for each condition and time point. We tested for differential expression between the WT and *lrp::kanR* strains separately for each condition and time point by using a Wald test on the genotype term of the simple model: transcript abundance ∼ genotype; here the *lrp::kanR* is the baseline condition. Transcripts that passed both an FDR corrected p-value of less than 0.05 and a genotype term magnitude of greater than 0.5 were considered as having a significant Lrp-dependent RNA expression change under that condition.

### Visualization of Lrp-dependent changes in Transcription

In order to visualize the Lrp-dependent changes in transcription data in a more intuitive manner, for all RNA expression bar graphs, we reported the following estimate of the log2 ratio of WT transcripts per million (TPM) over *lrp::kanR* TPM:

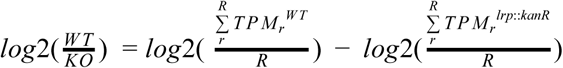

Where the total replicates *R* = 2 in all cases. To generate the error bars on all RNA expression bar plots, the log2(WT/KO) TPM was calculated as above for all 100 bootstrap replicates from kallisto, and a percentile based 95 % confidence interval from these bootstrap replicates was taken to be the lower and upper bounds of the ratio.

### Antibody development and testing

The monoclonal antibody used in these experiments was developed via a contract with NeoClone (Madison, WI). Using purified His-tagged Lrp, several rounds of potential antibodies were developed. The potential antibodies were tested for cross-reactivity with the known Lrp homologues AsnC and YbaO by ELISA at NeoClone. We used an *in vitro* DNA pull-down assay to ensure that the potential antibodies did not inhibit Lrp-DNA binding (Fig S6A). In addition, we tested the antibody for use in Western blotting (Fig S6B), and confirmed that the antibody did not bind the oligomerization interface by observing bands corresponding to Lrp octamers and hexadecamers in native Western blots.

### Filtering of genes into Lrp-dependent categories

For gene target filtering, we established four categories through a two-level filtering scheme (Fig. 2A). We first tested whether the gene had a Lrp-dependent change in RNA expression by comparing the target gene’s expression in WT and *lrp::kanR* strains using a Wald test as described above. We next asked if the gene had a high confidence Lrp binding site, as defined above, within the regulatory region (defined as 500 bp upstream and downstream from the annotated transcription start site (TSS; annotations from RegulonDB [18]). If multiple TSSs were annotated for a gene, the regulatory region included 500 bp upstream of the most distal TSS and 500 bp downstream of the most proximal TSS.

Using our high confidence Lrp binding regions, we then determined which regulatory regions fell within a high-confidence Lrp binding site; any regulatory region that overlapped with a high-confidence Lrp binding site was classified as bound by Lrp. Genes were thus categorized as either a direct target (RNA expression change and Lrp binding), an indirect target (RNA expression change but no Lrp binding), a NAP target (no RNA expression change but Lrp binding), or unconnected to Lrp (neither RNA expression change or Lrp binding).

For comparing enrichment of Lrp targets with σ factor targets, we used permutation tests as noted in the text, implemented using custom python scripts and 1000-10000 permutations. When testing for enrichment across several different σ factors, we corrected for multiple hypothesis testing using the statsmodels.sandbox.stats.multicomp.multipletests module using the Benjamini-Hochberg method [64,68].

All plots except where noted were created using ggplot2 [69] or Matplotlib [70].

## Acknowledgements/Funding

The authors wish to thank Prof. Robert Blumenthal for his advice and suggestions on this project. This research was supported by NIH R00 GM097033 (to PLF) and startup funds from the University of Michigan (to PLF). GMK is supported by the Cellular Biotechnology Training Program (T32-GM008353) and a one-time Rackham Research Grant from the University of Michigan. MBW is supported by a National Science Foundation Graduate Research Fellowship.

## Author Contributions

Planned experiments (GMK and PLF). Performed experiments (GMK). Performed sequencing analysis (MBW). Analyzed data (MBW, GMK and PLF). Prepared manuscript (MBW, GMK and PLF). Supervised the work (PLF).

